# Mechanistic insight into the oligomerisation of *Arabidopsis* CRY1 and its inhibition by BIC1

**DOI:** 10.1101/2025.10.06.680677

**Authors:** Alicia Just, Nils Niemann, Petra Gnau, Dennis Kock, Thomas Heimerl, Lars-Oliver Essen, Alfred Batschauer, Nina Morgner

**Affiliations:** Institute of Physical and Theoretical Chemistry, Johann Wolfgang Goethe University, Frankfurt/Main, Germany; Department of Biology, Philipps University Marburg, Marburg, Germany; Department of Chemistry, Philipps University Marburg, Marburg, Germany; SYNMIKRO Research Center and Department of Chemistry, Philipps University Marburg, Marburg, Germany

## Abstract

Cryptochromes (CRY) convert blue light signals into biological responses, however, the molecular processes underlying their activation are not fully understood. In this study, we uncover the molecular mechanism underlying the blue-light activation of *Arabidopsis* CRY1 using time-resolved native mass spectrometry combined with kinetic modelling – an approach that allows us to monitor light-driven complex formation with temporal and molecular resolution. We show that CRY1 activation follows a defined, reversible assembly pathway in which monomers rapidly form dimers that then assemble into tetramers. A quantitative two-step model captures the dynamic interplay between light-driven assembly and thermal disassembly. Strikingly, ATP accelerates tetramer formation and stabilises oligomers by tuning the underlying photochemistry of the FAD chromophore. In contrast, the Blue-light Inhibitor of Cryptochromes 1 (BIC1) acts as a potent antagonist. We found that BIC1 binds to CRY1 even in the dark, with a significant increase in binding strength under blue-light conditions. BIC1 not only blocks CRY1 oligomerisation, but also actively dismantles pre-assembled tetramers. This disassembly process is light-independent and occurs regardless of CRY1’s redox state. Together, these findings reveal a finely balanced regulatory system in which ATP and BIC1 act as opposing regulators to control CRY1 activation. This work provides a kinetic and mechanistic framework for reversible cryptochrome signalling and highlights how blue light responses can be precisely modulated at the molecular level.

## Introduction

Plants rely on light not only as an energy source but also as a critical environmental signal that guides nearly every aspect of their growth and development^1–3^. To perceive and translate light information across the spectrum, plants have evolved a sophisticated network of photoreceptors, including phytochromes, phototropins, the UV-B photoreceptor UVR8, and cryptochromes (CRYs)^1,4–6^. Among these, CRYs are evolutionarily conserved flavoproteins that absorb blue and ultraviolet-A light via a flavin adenine dinucleotide (FAD) chromophore, which is bound non-covalently within the photolyase homology region (PHR) domain^7–10^. The PHR domain is complemented by the cryptochrome C-terminal extension (CCE) that is much less conserved and varies in length and sequence among cryptochromes^11^. In *Arabidopsis thaliana*, two major cryptochromes, CRY1 and CRY2, operate in partially overlapping but distinct physiological contexts^12^. CRY1 predominantly controls the early seedling stage, where it suppresses hypocotyl elongation, whereas CRY2 regulates flowering time by acting as a photoperiod sensor^12–16^. Activation of plant CRYs is initiated by light-driven photoreduction of FAD^17,18^, which induces conformational changes that promote oligomerisation^19–22^ and binding to downstream signalling partners^23^. Although CRY1 and CRY2 are structurally related^24^, their functional specialization suggests that they may differ in their activation dynamics, signalling interactions, and mechanisms of regulation. Structural studies have demonstrated that both CRY1 and CRY2 undergo blue light–induced oligomerisation^13,15–17^, and in the case of CRY2, tetrameric assemblies have been linked to signalling activity^25^. CRY1 likewise forms light-induced oligomers, as revealed by structural analyses^26^, yet the dynamics and regulation of this process are poorly understood. While oligomerisation is mediated by the PHR domain, the CCEs are engaged in liquid–liquid phase separation (LLPS)^11^. To distinguish specific oligomerisation from the formation of dynamic size-variable condensates, *in vitro* studies of the isolated PHR domain are essential to uncover the intrinsic mechanism of cryptochrome oligomerisation without the interfering influence of LLPS. In particular, the molecular mechanism underlying CRY1 oligomerisation as well as the influence of cellular cofactors or inhibitory proteins on this process, remains poorly understood.

A major class of negative regulators of CRYs are the Blue-light Inhibitors of Cryptochromes (BICs)^27^. BICs interact with CRY2 in its PHR domain and efficiently block tetramerization, thereby suppressing signalling^21,24^. CRY1 has also been identified as a target of BIC1^21,25–27^, but the mechanistic details of this inhibition are not resolved. Previous studies demonstrated that BIC1 may associate with CRY1 in both dark and light conditions^28^, raising the question whether BIC1 primarily prevents oligomer formation or whether it can actively dismantle pre-assembled CRY1 complexes.

Another layer of potential regulation arises from cellular metabolites^29–31^. ATP is known to bind the PHR domain of plant cryptochromes^17,18^ and a variety of studies reported, that ATP accelerates FAD photoreduction and stabilizes reduced states^30,32–34^. However, whether ATP directly influences CRY1 oligomerisation dynamics has remained unclear.

Building on our previous work, where we demonstrated time-resolved investigation of light-induced conformational changes in the monomeric animal-like cryptochrome from *Chlamydomonas reinhardtii* (*Cr*aCRY)^35^, we now take one step further and explore the oligomerisation process that follows photoactivation in CRY1 from *Arabidopsis thaliana*. We combined different native mass spectrometry techniques, absorption spectroscopy and negative-stain transmission electron microscopy, to unravel the light-induced assembly and regulation of *Arabidopsis* CRY1 PHR domain *in vitro*. We show that CRY1 oligomerisation proceeds via a sequential monomer → dimer → tetramer pathway that can be quantitatively described by a reversible two-step kinetic model. ATP accelerates this process, boosting the oligomerisation rates and stabilizing the oligomeric states. Furthermore, we demonstrate that BIC1 binds to CRY1 both in darkness and under blue light illumination, with significantly different affinities. Importantly, BIC1 not only prevents tetramer assembly but also disassembles preformed CRY1 oligomers, a property that extends to the constitutively tetrameric CRY1^L407F^ mutant^36,37^. Finally, we show that BIC1 inhibition does not interfere with photoreduction of the FAD chromophore, indicating that BIC1 regulates CRY1 through direct competition with CRY1-CRY1 contacts rather than through modulation of chromophore photochemistry.

Together, our findings establish a quantitative kinetic framework for CRY1 oligomerisation and reveal dual regulatory inputs: metabolic tuning by ATP and active disassembly by BIC1. These results uncover fundamental principles of CRY1 regulation that distinguish it from CRY2 and highlight new mechanisms by which plant cells integrate metabolic state and the effect of inhibitory proteins to fine-tune cryptochrome signalling.

## Results

### Oligomerisation mechanism of *Arabidopsis* CRY1

To gain mechanistic insight into *Arabidopsis* CRY1 oligomerisation, we investigated the oligomeric states of the recombinant CRY1 PHR domain (CRY1-PHR), thereafter named CRY1, *in vitro* by nano electrospray ionization mass spectrometry (nESI-MS). Consistent with previous reports^26^ CRY1 exists predominantly as a monomer in the dark (m/z = 3500-4800, Fig. 1a) with only a small fraction present as dimers (m/z = 5000-5700). Exposure to blue light promotes its oligomerisation into a tetrameric form (m/z = 6000-8500) (Fig. 1b).

**Fig. 1:**
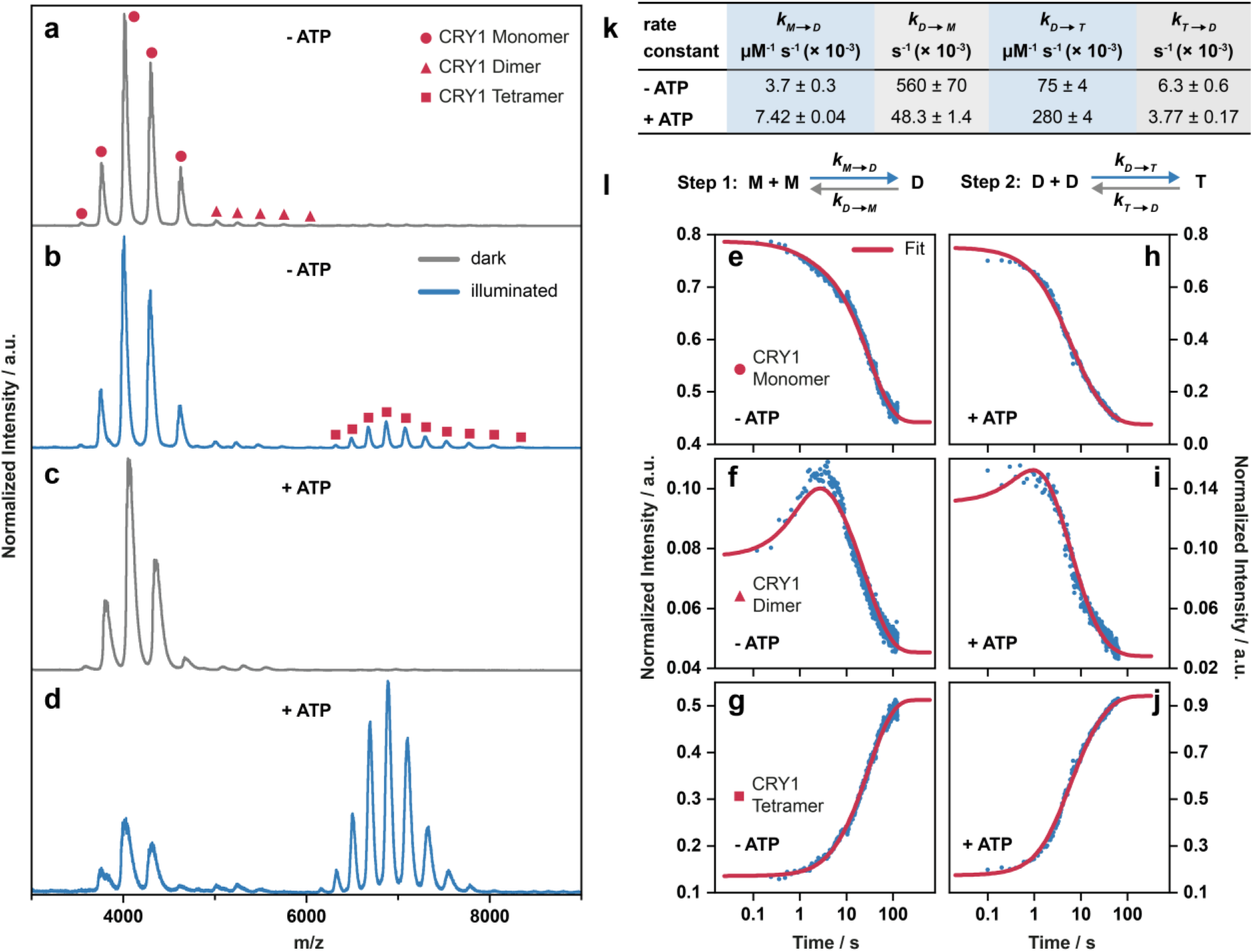
Oligomerisation kinetics of CRY1 under blue light and the influence of ATP. nESI-MS spectrum of CRY1 in the dark (grey line) (**a, c**) and under illumination with blue light (blue line) (**b, d**) in absence **(a, b**) and presence of 100 µM ATP (**c, d**). Exposure to blue light leads to oligomerisation of CRY1, and the addition of ATP promotes this process. Time-resolved mass spectrometry measurements of the CRY1 monomer (**e**), dimer (**f**) and tetramer (**g**) in absence and presence of ATP (**h-j**) and the fit (pink line) obtained from the two-step reversible kinetic model (**l**). **k,** Association (highlighted in blue) and dissociation (highlighted in grey) rates obtained from kinetic modelling in presence and absence of 100 µM ATP.

To elucidate the underlying mechanism of this light-induced oligomerisation, we next examined whether tetramer assembly occurs exclusively via dimer intermediates or involves trimeric species as transitional state. To address this, we performed time-resolved mass spectrometry at room temperature, allowing us to detect and quantify transient oligomeric species under blue light conditions. The time-resolved data revealed a well-defined kinetic progression of CRY1 oligomerisation (Fig. 1e-g). In the initial dark-adapted state, the cryptochrome was predominantly monomeric (Fig. 1a, e). Upon exposure to blue light, we observed a rapid decrease in monomer abundance (Fig. 1e), accompanied by the appearance of dimers (Fig. 1f). Tetramer formation was observed after a short delay, consistent with a stepwise assembly pathway (Fig. 1g). Dimer levels peaked after approximately 3 seconds (Fig. 1f), while tetramer accumulation progressed more slowly and reached a stationary plateau at 120 seconds (Fig. 1g). Notably, the monomeric species did not fully disappear (Fig. 1b, e), suggesting that oligomerisation is opposed by a concurrent thermal dissociation process. This observation shows that even under sustained light activation, reversible oligomer assembly and disassembly are in kinetic competition.

To quantitatively describe the oligomerisation mechanism, we propose a reversible kinetic model comprising two steps as shown in Fig. 1l. This kinetic model assumes sequential assembly, from monomer via dimer to tetramer (M→D→T), with no direct conversion from monomer to tetramer and no trimeric species. Fitting our data with this model (Fig. 1e-g, Extended Data Fig. 2a) yielded association and dissociation rate constants listed in Fig. 1k. The model accurately described the experimental time courses of all three oligomeric states at room temperature. The relatively low value of k_M→D_ = (3.7 ± 0.3) × 10^-3^ µM^-1^ s^-1^ indicates slow dimerization, while the larger association rate k_D→T_ = (75 ± 4) × 10^-3^ µM^-1^ s^-^1 reflects a more efficient tetramerization step once dimers are present. The dissociation rates further reveal distinct stabilities: the dimer dissociates rapidly with k_D→M_ = (560 ± 70) × 10^-3^ s^-1^, whereas the tetramer is more stable (k_T→D_ = (6.3 ± 0.6) × 10^-3^ s^-1^). Together these rate constants collectively explain the observed accumulation of tetramers and the persistence of monomers in a dynamic light-induced stationary state. These results demonstrate that the experimentally observed oligomerisation dynamics of CRY1 can be effectively described by a two-step reversible reaction model and provide quantitative insight into the kinetics of light-induced assembly.

### ATP boosts oligomer formation

Given previous reports that ATP influences the FAD photoreduction kinetics of cryptochromes^30,32–34^, we next investigated whether ATP also affects the light-induced oligomerisation of CRY1. Under identical conditions in presence and absence of ATP, we observed a pronounced ATP-dependent enhancement of tetramer formation (Fig. 1c, d). In the presence of 100 µM ATP, CRY1 assembled into tetramers more efficiently and reached markedly higher tetramer levels upon blue light exposure (Fig. 1d, Extended Data Fig. 2b). Moreover, the rate of tetramer oligomerisation almost quadrupled, while the subsequent thermal dissociation in darkness was slowed approximately twofold for the tetramer and even more than tenfold for the dimer (Fig. 1h-k). To further assess the concentration dependence of this effect, we also tested a concentration of 200 µM ATP. Under these conditions, we detected a substantial increase in the dimeric CRY1 population already in the dark-adapted state compared to samples without ATP or in presence of 100 µM ATP (Extended Data Fig. 1a-c). This observation suggests that higher ATP concentrations promote or stabilize dimer formation even in the absence of blue light. These findings show that ATP enhances and stabilizes dimer formation, which we observed to be the rate limiting step for tetramer formation under blue light conditions, revealing a broader modulatory role of ATP in tuning cryptochrome oligomerisation dynamics. Absorption spectra also show the boosting effect of ATP on the kinetics of FAD photoreduction, as has already been shown in previous studies^30,32–34^. In our measurements the addition of ATP accelerated the photoreduction of FAD_ox_ to FADH° by approximately 12%, while slowing down the thermal decay of FADH° back to FAD_ox_ by nearly 60% (Extended Data Fig. 1d-h).

### BIC1 binds CRY1 in both darkness and under blue light

While BIC1 is known to bind CRY1, it remains unresolved whether this interaction blocks CRY1 oligomerisation or dismantles existing assemblies. We therefore directly investigated the mechanism of CRY1 inhibition by BIC1 using Laser-Induced Liquid Bead Ion Desorption mass spectrometry (LILBID-MS)^38^. In line with the nESI-MS data, LILBID-MS analysis further confirmed that CRY1 exists predominantly as a monomer in the dark (∼ 59 kDa), with only a minor dimeric population (∼ 120 kDa) detectable (Fig. 2a). Blue-light illumination shifted the oligomeric distribution rapidly toward tetrameric assemblies (∼ 240 kDa), underscoring light-dependent higher-order organization of CRY1 (Fig. 2b). A small trimeric CRY1 population was also detected, arising from partial dissociation of the tetramer induced by the energy input during LILBID-MS analysis (Extended Data Fig. 3a). Upon addition of BIC1 (∼ 20 kDa) in both the dark and under blue light conditions, we observed a discrete charge distribution corresponding to the mass of CRY1-BIC1 heterodimer (∼ 79 kDa), indicating direct binding between the two proteins (Fig. 2c, d). While CRY1 exhibited reduced binding to BIC1 in the dark (Fig. 2c), we observed a significant increase in CRY1-BIC1 heterodimer formation under blue light conditions (Fig. 2d). Notably, in light-exposed samples, the CRY1 tetramer signal was absent, suggesting that BIC1 binding prevents oligomer formation. To quantify the interaction strength of CRY1-BIC1 heterodimer in the dark and under blue light conditions, we employed quantitative LILBID-MS (qLILBID-MS), a label-free technique capable of measuring dissociation constants under native-like conditions^39,40^. In the dark, BIC1 bound CRY1 with moderate affinity (K_D_ ≈ 1.2 µM) (Fig. 2g), whereas under blue light, the interaction became more stable (K_D_ ≈ 0.13 µM) (Fig. 2f). This ten-fold increase in affinity suggests that light-induced conformational changes in CRY1 enhance its recognition by BIC1. This demonstrates that BIC1 binds CRY1 in both its inactive and active state but with a strong preference for the light-activated form, consistent with a mechanism in which BIC1 selectively inhibits photoactivated CRY1.

**Fig. 2:**
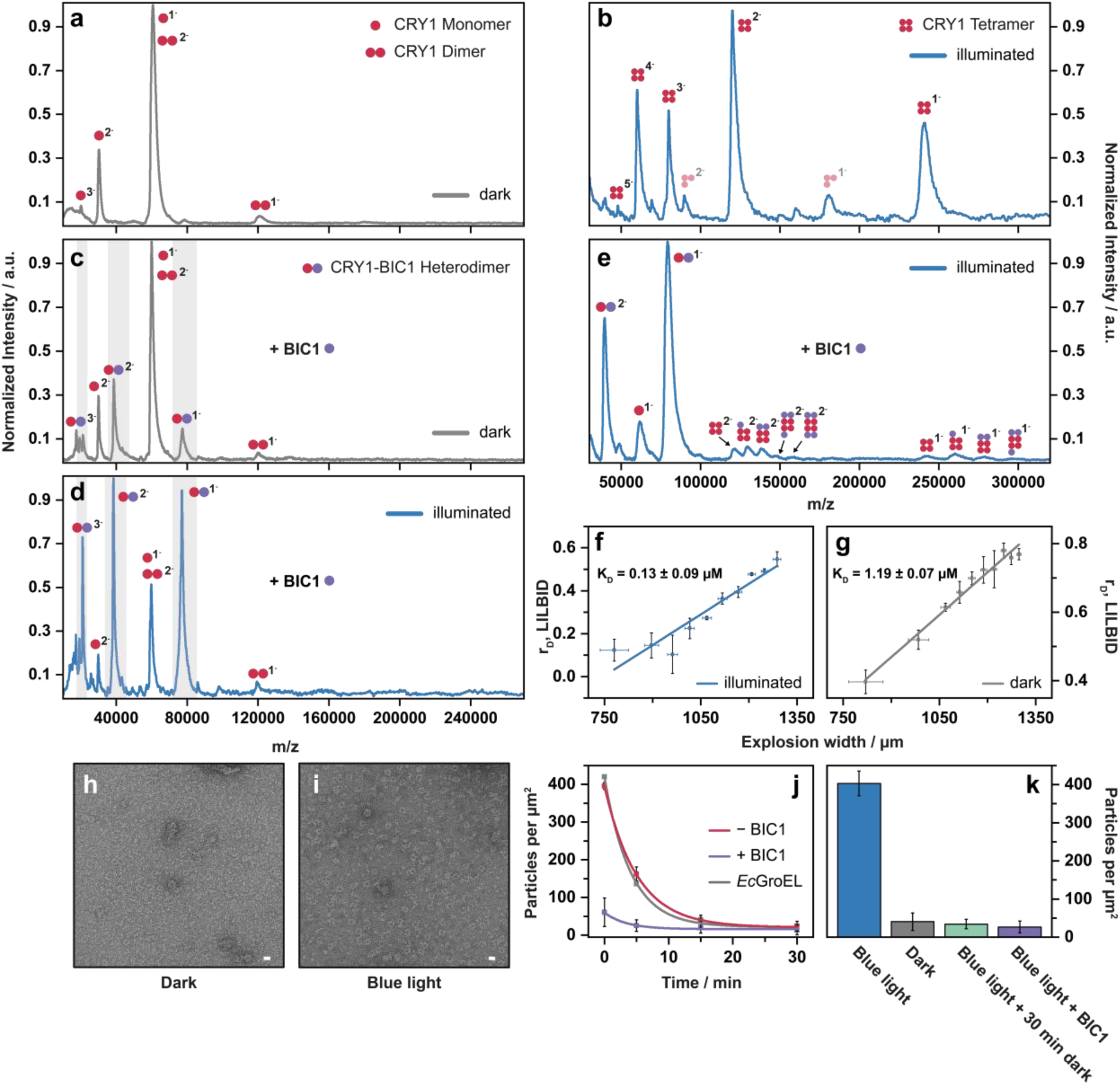
BIC1 inhibits CRY1 oligomerisation in both dark and blue light conditions with higher affinity of BIC1 to CRY1 under blue light conditions. LILBID-MS spectrum of CRY1 shows a monomer (pink dot) (**a**) in darkness, which forms tetramers upon illumination with blue light (**b**). CRY1 trimers are caused by laser dissociation. **c,** Incubation of CRY1 monomer and BIC1 (purple dot) in an equimolar ratio in darkness results in a mixture of CRY1-BIC1 heterodimer and CRY1 monomer. **d,** Upon blue light exposure, the equilibrium shifts towards a higher proportion of the CRY1-BIC1 heterodimer relative to the CRY1 monomer (highlighted in grey). **e,** Under blue light conditions, addition of BIC1 to the CRY1 tetramer results in binding to each protomer, thereby dissociating the tetramer into CRY1-BIC1 heterodimers. f, qLILBID-MS measurements reveal that the CRY1-BIC1 heterodimer exhibits a ten-fold higher affinity under blue light compared to its dark-state counterpart (**g**). TEM images of CRY1 particles visualised by negative staining of probes kept in darkness (**h**), or treated with blue light (450 nm, 20 min, 115 µmol m ^2^ s^-1^) (**i**) showing light-induced formation of doughnut-like ring structures of CRY1. Scale bars 10 nm. **j**, Kinetics of CRY1 monomerization (n = 3 ± SD) after transfer of samples from blue light to darkness in absence and presence of BIC1 or *E. coli* GroEL as negative control. **k**, Effect of BIC1 on kinetics of decrease in number of CRY1 particles showing monomerization of CRY1 (n = 3 ± SD). Blue light (465 nm, fluence rate 115 µmol m^-2^ s^-1^) treatment was for 20 min.

### BIC1 reverses CRY1 oligomerisation

Given that CRY1 tetramerizes upon blue light exposure (Fig. 2b), and that BIC1 binds with high affinity under blue light (Fig. 2d), the question arises whether BIC1 merely blocks tetramer formation or can even actively dismantle pre-existing CRY1 oligomers as it was shown for CRY2 and BICs^21,24^. Therefore, we added BIC1 to pre-illuminated, tetrameric CRY1 under constant illumination with blue light. The mass spectra indicate that BIC1 binds sequentially to individual CRY1 protomers within the tetramer (∼ 260-320 kDa) (Fig. 2e), displacing them from the tetrameric complex and a formation of a dominant BIC1–CRY1 heterodimer (∼ 79 kDa). These data suggest that BIC1 not only prevents tetramer formation but also actively dismantles existing CRY1 tetramers by occupying all subunits.

Negative-stain transmission electron microscopy (TEM) corroborates the mass spectrometry data. In blue light-treated samples, CRY1 formed doughnut-like ring structures that were absent in dark-adapted preparations (Fig. 2h, i), consistent with the formation of tetrameric assemblies. The addition of BIC1 markedly reduced the abundance of these oligomeric particles. Even immediately after 30 minutes of illumination, CRY1 samples incubated with BIC1 displayed few ring-like structures (Fig. 2k). Moreover, following transfer to darkness, CRY1 oligomers disassembled significantly faster in the presence of BIC1, with near-complete monomerization observed at the earliest time point (Fig. 2j).

### BIC1 disassembles CRY1 tetramers independently of light activation

To examine whether BIC1-mediated inhibition of CRY1 tetramers requires light activation, we analyzed the CRY1^L407F^ mutant, which has been observed to be hyperactive and results in a constitutive photomorphogenic phenotype^36,37^. Analytical size exclusion chromatography (SEC) revealed, that CRY1^L407F^ shows already a tetrameric retention time in darkness compared to the wild-type CRY1 (Fig. 3d). LILBID-MS confirmed the presence of tetramers for CRY1^L407F^ in darkness (Fig. 3a) and absorption spectroscopy has shown that the leucine to phenylalanine replacement at position 407 does not impair photoreduction of the chromophore (Fig. 3h).

**Fig. 3:**
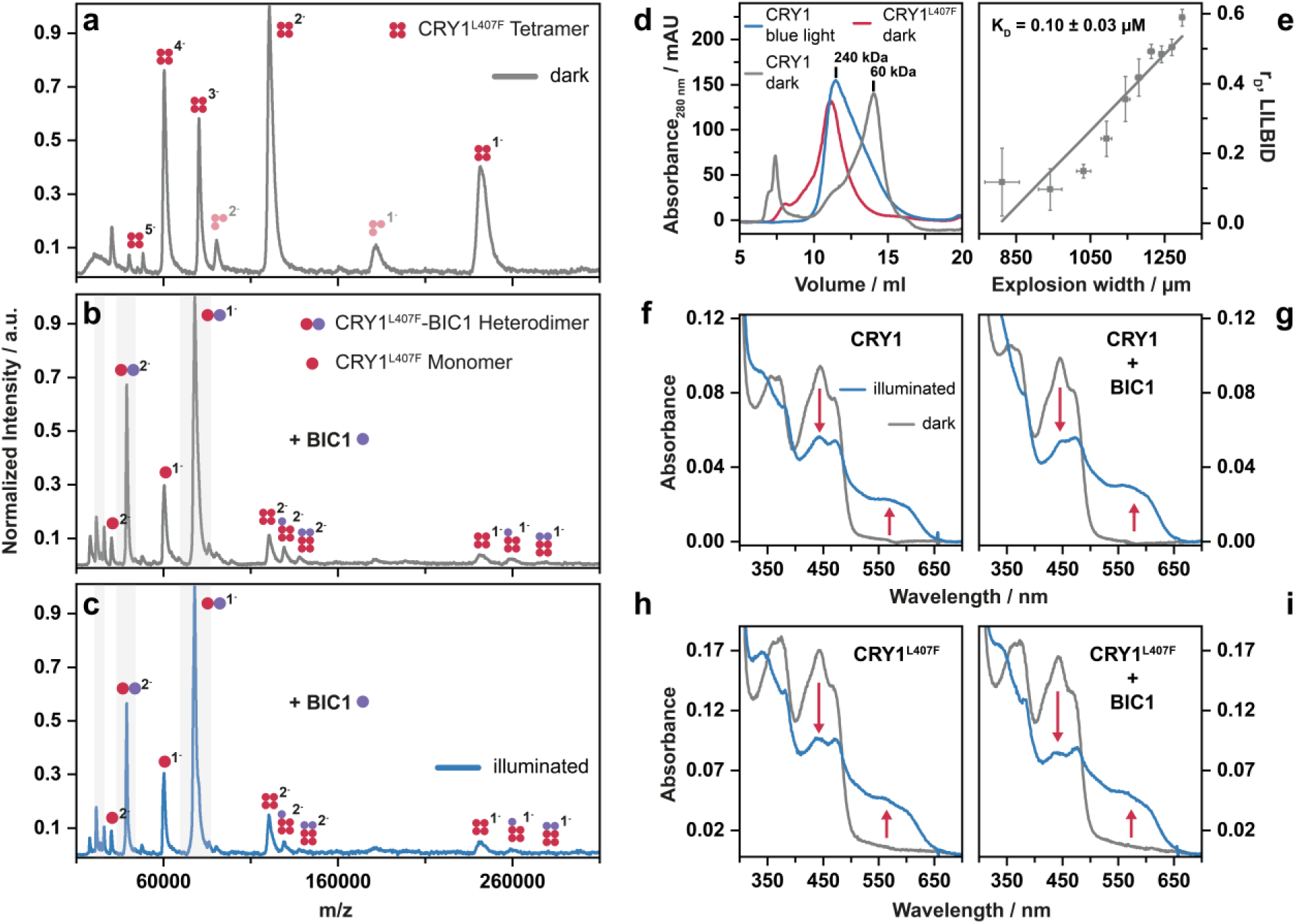
BIC1 disassembles constitutively active CRY1 tetramers and does not inhibit photoreduction of the FAD cofactor. **a**, LILBID-MS spectrum of the constitutively active CRY1^L407F^ mutant shows a tetramer in darkness. CRY1 trimers are caused by laser dissociation. **b**, Addition of BIC1 (purple dot) to CRY1^L407F^ in an equimolar ratio in darkness results in binding to each protomer, thereby dissociating the CRY1^L407F^ tetramer into CRY1^L407F^-BIC1 heterodimer and monomeric CRY1^L407F^. **c**, Exposure to blue light does not shift the equilibrium toward the CRY1^L407F^-BIC1 heterodimer, as has been shown for wild-type CRY1. **d**, SEC elution profiles of wild-type CRY1 kept in darkness or irradiated with blue light, and of hyperactive CRY1^L407F^ mutant kept in darkness. **e**, qLILBID-MS measurement of the CRY1^L407F^-BIC1 heterodimer under dark conditions revealed that BIC1 binds CRY1^L407F^ with high affinity. **f**, Absorption spectra of wild-type CRY1 before (grey line) and after (blue line) blue light exposure indicate FAD photoreduction, with a decrease in FADox peak at 450 nm, and the emergence of a broad band at 550-600 nm, characteristic for the FADH° semiquinone. **g**, Addition of BIC1 to wild-type CRY1 does not inhibit the photoreduction of CRY’s FAD chromophore. **h**, Absorption spectra of the CRY1^L407F^ mutant indicates that chromophore photoreduction remains unaffected by the mutation. **i**, Photoreduction of the FAD chromophore in the CRY1^L407F^ mutant remains unaffected by BIC1 addition, as observed for wild-type CRY1.

Upon addition of BIC1, the CRY1^L407F^ tetramers were efficiently disassembled into BIC1–CRY1^L407F^ heterodimers (Fig. 3b, Extended Data Fig. 3b). In contrast to the wild-type CRY1 (Fig. 2c, d), exposure to blue light did not further enhance the formation of the CRY1^L407F^-BIC1 heterodimer (Fig. 3c) or have any other significant effect on the observed complexes. qLILBID measurements revealed that BIC1 binds CRY1^L407F^ independently of illumination with high affinity (K_D_ = 0.10 ± 0.03 µM), comparable to the affinity of BIC1 for light-activated wild-type CRY1 (K_D_ = 0.13 ± 0.09 µM). This result demonstrates that the L407F mutation induces a conformational change in CRY1 that mimics its light-activated state with respect to BIC1 recognition.

Collectively, these findings suggest that BIC1 disrupts CRY1 oligomerisation by binding individual CRY1 protomers with higher affinity than the protomer-protomer interactions, effectively outcompeting CRY1-CRY1 contacts regardless of the photoreduction state of the chromophore.

### BIC1 does not inhibit photoreduction of the CRY1 FAD cofactor

The remaining open question is whether BIC1 suppresses CRY1 oligomerisation by interfering with its photoactivation, as previously demonstrated for CRY2 inhibition by BIC2^24^. To address this, we examined the photoreduction state of the FAD chromophore in the presence and absence of BIC1. Absorption spectra of purified CRY1 recorded before and after blue light exposure showed the expected spectral changes associated with FAD photoreduction (Fig. 3f, Extended Data Fig. 1d, e). In the dark-adapted state, CRY1 displayed absorption maxima characteristic of oxidized flavin (FAD_ox_) at 450 nm and a shoulder near 480 nm. Upon blue light illumination, these peaks decreased in intensity, accompanied by the appearance of a broad band between 550-600 nm, consistent with the formation of the neutral semiquinone radical (FADH°)^17,18,41^. These spectral changes are characteristic for the light-driven redox transition of the CRY1 chromophore. Identical spectral features were observed when CRY1 or CRY1^L407F^ were incubated with BIC1 under the same conditions (Fig. 3g, i, Extended Data Fig. 4a-c), indicating that BIC1 does not impair the light-induced redox transition of the FAD_ox_ to FADH°. This suggests that BIC1 does not block the initial photoactivation step of CRY1. Instead, it acts downstream of photoreduction, targeting the light-induced oligomeric states of CRY1.

## Discussion

Light perception in plants is a dynamic process influenced by multiple molecular factors. Our kinetic analysis of CRY1 reveals that oligomerisation proceeds through discrete assembly steps and is modulated by both metabolic signals and inhibitory interactions. These findings suggest that CRY1 activity can be finely adjusted in response to changing conditions, enabling a more nuanced regulation of light signalling.

Our time-resolved native mass spectrometry data establish a clear kinetic pathway for CRY1 oligomerisation (Fig. 4). The sequential monomer-to-dimer-to-tetramer assembly we observed is consistent with a two-step reversible kinetic model (Fig. 1l), identifying dimer formation as the rate-limiting step in CRY1 activation. In comparison, FAD photoreduction occurs within microseconds^42^, much faster than oligomerisation, indicating that the light-induced activation of the flavin cofactor does not contribute any significant kinetic delay. The relative stabilities of the oligomers differ substantially, with dimers being highly transient while tetramers display significantly increased stability. These light dependent equilibria, suggests that CRY1 signalling is a highly dynamic process rather than a simple on-off switch. To directly compare our *in vitro* oligomerisation kinetics with *in vivo* observations made by Liu *et al*. (2020)^22^, we fitted the tetramer formation data using a single-exponential model and determined a half-life of approximately t_1/2_ = 18 s (Extended Data Fig. 2c). *In vivo*, the corresponding half-life at a fluence rate of 30 µmol m^−2^ s^−1^ is about five times longer, reflecting additional cellular constraints such as regulatory interactions (e. g. BIC1 inhibition), metabolites, different reducing environment, temperature and, of course, different protein concentrations. Despite these differences, the timescales remain within the same order of magnitude, underscoring the physiological relevance of the oligomerisation process captured by time-resolved mass spectrometry. Importantly, our *in vitro* approach allows disentangling the influence of individual factors that cannot easily be resolved in the cellular context, thereby complementing *in vivo* measurements. Beyond this biological relevance, such a detailed kinetic analysis also opens opportunities for engineering CRY1-based optogenetic tools with tuneable temporal precision, complementing existing CRY2-based optogenetic systems^43–46^.

**Fig.4:**
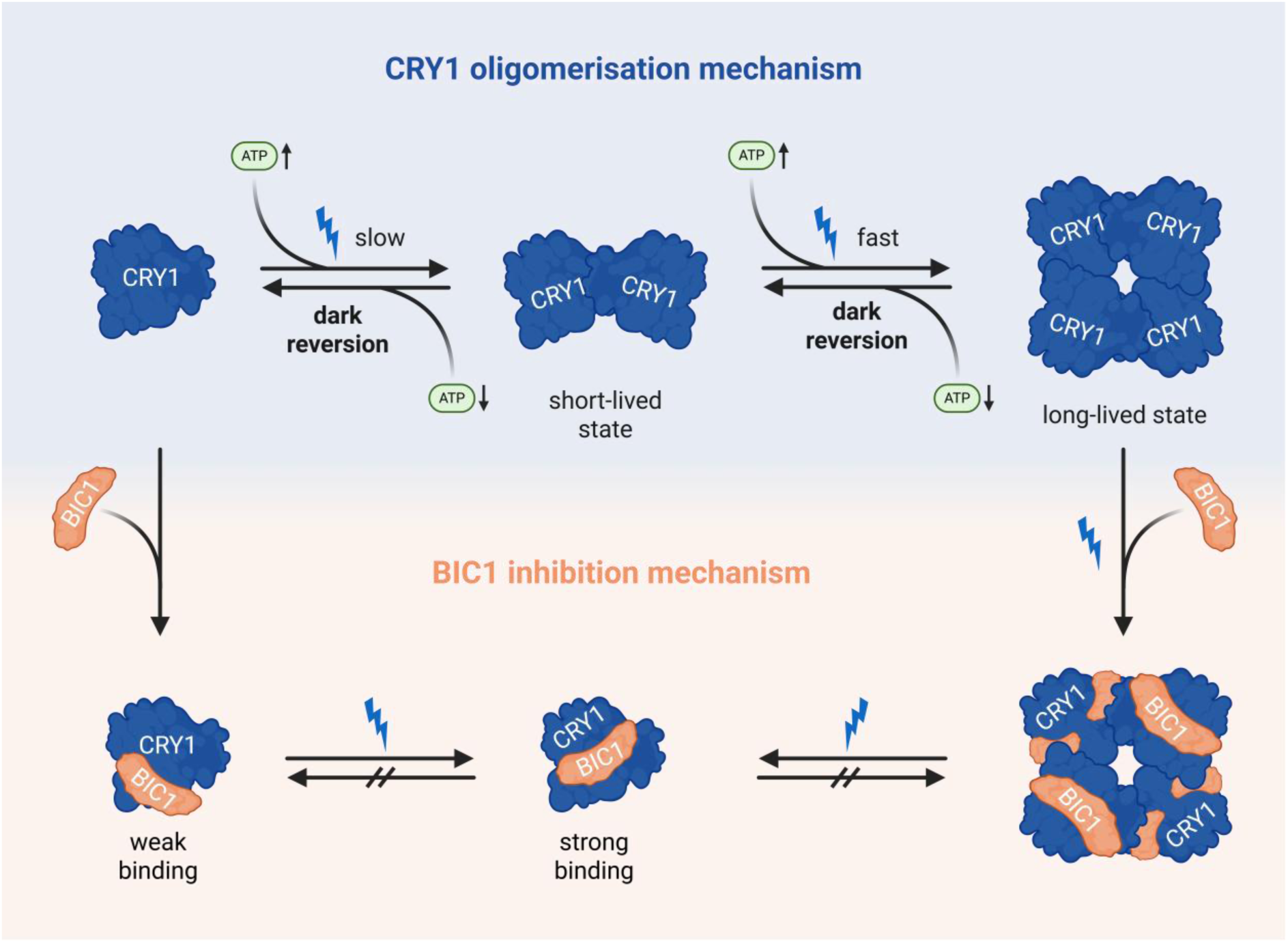
Schematic model of CRY1 oligomerisation dynamics, ATP modulation, and BIC1-mediated inhibition. Upon blue light exposure, CRY1 transitions from a monomeric to a tetrameric state via a short-lived dimeric intermediate. This oligomerisation follows a sequential, reversible two-step mechanism. ATP enhances the association rates for dimer and tetramer formation, while slowing down the dissociation rates of these oligomers. BIC1 binds CRY1 in both dark and light conditions, with significantly higher affinity for the light-activated conformation. This interaction prevents oligomer formation and actively disassembles pre-formed tetramers by binding to individual CRY1 protomers even under continuous blue-light illumination.

In addition to intrinsic kinetic insights, our findings highlight how cellular metabolites can modulate this process. We show that ATP dramatically enhances CRY1 oligomerisation upon illumination (Fig. 4). Increasing ATP concentrations significantly enhance the formation of CRY1 dimer (Extended Data Fig. 1a-c) which we showed to be the rate limiting step in oligomer assembly. Then tetramer formation proceeds rapidly, resulting in more stable higher-order oligomers.

This is particularly intriguing given that ATP is not only a fundamental energy currency but has also been reported to bind the PHR domain of plant cryptochromes^21,28^ and affect FAD photoreduction kinetics^30,32–34^. Our data support the notion that ATP acts as a cofactor, stabilizing the oligomeric state and promoting the conformational changes required for stable oligomer formation. This mechanism suggests a novel link between the cellular energy status and CRY1 activity, allowing a plant to modulate its light response in accordance with its metabolic state. For instance, it is conceivable that under conditions of high energy availability, CRY1 signalling may be sensitized to support energy-intensive processes such as development and growth.

Our data further elucidate the inhibitory role of BIC1, revealing an elegant mechanism of regulation (Fig. 4). We demonstrate that BIC1 binds to CRY1 in both dark and light conditions (Fig. 2c, d), but with an enhanced affinity for the light-activated form (Fig. 2f, g). This light-dependent increase in binding affinity indicates that light-induced conformational changes in CRY1 significantly promote its interaction with BIC1. This light-dependent increase in affinity is a crucial finding, as it provides a mechanism for the selective inhibition of active CRY1. Moreover, we provide clear evidence that BIC1 is an active disassembler of CRY1 oligomers, rather than just a passive competitor (Fig. 2e). Given that the constitutively active CRY1^L407F^ mutant, which mimics the tetrameric light-activated state, shows a high affinity for BIC1 (Fig. 3e), strongly suggests that BIC1 recognizes a specific conformational state of CRY1, independent of the FAD redox state. Our result that BIC1 does not interfere with FAD photoreduction (Fig. 3g, i) differs from recent reports on the CRY2-BIC2 interaction^24^, suggesting that the inhibitory mechanisms are distinct for CRY1 and CRY2. This supports the idea that while cryptochromes share a conserved photochemical activation step, their downstream regulation can vary, allowing for functional specificity.

In summary, our work establishes a quantitative kinetic model for CRY1 oligomerisation. By showing that ATP acts as an oligomerisation accelerator and BIC1 as an active disassembler, we provide a more detailed and nuanced understanding of how CRY1 activity is regulated beyond simple light perception. These insights not only shed light on the functional specialization of CRY1 but also highlight a broader principle of plant signalling, where a photoreceptor’s output is shaped by a complex interplay between light, metabolites, and dedicated regulatory proteins.

## Methods

### Cloning of constructs

The expression vectors encoding the PHR domain (residues 1–509) of *Arabidopsis thaliana* CRY1 and the CRY1^L407F^ mutant, both cloned into the pACYCDuet-1 vector have been described previously^26^. To prevent disulfide bond-mediated dimerization observed in the *Arabidopsis thaliana* wild-type BIC1 protein (Extended Data Fig. 5a, b), a synthetic gene encoding a BIC1 variant carrying three cysteine-to-alanine substitutions (C99A, C105A, C136A) was generated. This triple mutant, named BIC1 throughout the text, was synthesized and cloned into the pET28a(+) vector (*BioCat GmbH*). Recombinant proteins carried an N-terminal His₆-tag.

### Overproduction and purification of recombinant proteins

For overexpression of CRY1 and the CRY1^L407F^ mutant as well as BIC1^3CA^, competent *E. coli* BL21 Star (DE3) cells (*Thermo Fisher*) were used. For CRY1 and CRY1^L407F^ overproduction, 20 mL overnight cultures were inoculated into 2 L LB medium supplemented with 34 µg/mL chloramphenicol in 5 L baffled flasks. Cultures were incubated at 37 °C with shaking (110 rpm) until an OD_600 nm_ of ∼1.0 was reached. Cells were placed on ice for 30 min, and expression was induced with 0.25 mM IPTG. All further steps were carried out under red light. Cultures were incubated for 3 days at 16 °C. Cells were harvested by centrifugation (5,330g, 15 min, 4 °C) and resuspended in lysis buffer (50 mM MOPS, pH 7.4, 300 mM NaCl, 10% glycerol, 1% Triton X-100). Lysozyme (0.2 mg/mL), PMSF (0.4 mM) and DNase I were added, and the suspensions were incubated for 10 min at room temperature before disruption using a French press. Lysates were clarified by centrifugation (39,000g, 30 min) and filtered. The supernatant was applied onto a 1 mL Protino Ni-NTA column (*Macherey-Nagel*), followed by sequential washing steps with lysis buffer, buffer containing 3% Triton X-100, and lysis buffer without TX-100 until a stable UV baseline was reached. Non-specifically bound proteins were removed with 10% elution buffer (50 mM MOPS, pH 7.4, 300 mM NaCl, 10% glycerol, 250 mM imidazole). Recombinant CRYs were eluted with 100% elution buffer. CRY1 concentrations were kept below 1 mg/mL to prevent oligomerisation. Final purification was performed by size-exclusion chromatography (Superdex 200 pg 16/60, *Cytiva*).

BIC1 purification followed a similar protocol. Expression was induced with 0.5 mM IPTG, and cultures were incubated for 18 h at 25 °C. Cells were harvested, washed with PBS, and resuspended in lysis buffer (50 mM HEPES, pH 7.0, 500 mM NaCl, 10% glycerol). Cell disruption was performed as described above, using a protease inhibitor mix (*Carl Roth, Cat. no. 3760.1*) in place of PMSF. The soluble fraction was loaded onto a Ni-NTA column and washed with protein buffer (50 mM HEPES, pH 7.0, 300 mM NaCl, 10% glycerol), followed by a wash containing 3% Triton X-100 and protein buffer. Additional washes were performed with 10% and 20% elution buffer (50 mM HEPES, pH 7.0, 300 mM NaCl, 250 mM imidazole, 10% glycerol). BIC1 was eluted with 100% elution buffer, concentrated, and further purified via size-exclusion chromatography (Superdex 75 pg 16/60, *Cytiva*).

### Size exclusion chromatography (SEC)

For analysis of blue light-dependent oligomerisation, samples containing 30 μM CRY1 and 10 mM ß-mercaptoethanol in 100 µl volumes were separated on a Superdex 200 10/300 GL column (*GE Healthcare*) equilibrated at 10 °C in buffer D containing 20 mM ß-mercaptoethanol. CRY1 was pre-illuminated with blue light (465 ± 5 nm, 115 µmol m^−2^ s^−1^) for 20 min. Three blue light LEDs were used to illuminate the column resin. Dark controls were kept in red safe light. Calibration of the column was done using gel-filtration standards (*Sigma Aldrich*).

### Photoreduction and dark reversion of CRY1

Absorbance changes of bound flavin were measured by UV-Vis spectroscopy (*Specord 600, Analytik Jena*). Before photoreduction, the CRY1 solution was centrifuged (10 min, 16873 g, 4 °C) to pellet and remove aggregated protein. A protein solution of 10 µM CRY1 was prepared, and dithiothreitol (DTT) added to a final concentration of 10 mM. For photoreduction, samples were irradiated with a 455 nm LED (M455L2, *Thorlabs*) at 100 µmol m^−2^ s^−1^. At defined time points, the absorption between 200 nm – 900 nm was recorded. The dark conversion was monitored with the same photoreduced sample by leaving it in the dark until full recovery and absorbance recording. During all steps, cuvettes were chilled at 20 °C.

### Nano Electrospray ionization mass spectrometry

nESI-MS was carried out using a Synapt G2-S instrument (*Waters Corporation, Wilmslow, UK*) equipped with a high-mass quadrupole modification. Nano electrospray emitters are produced in-house by pulling borosilicate glass capillaries with a Flaming/Brown micropipette puller (P-1000, *Sutter Instrument Co.*) and subsequently coated with a gold layer via sputtering. CRY1 (15 µM) was analyzed at room temperature in positive ion mode under a capillary voltage of 1.1 kV. The ion source was operated with a cone voltage of 150 V, an offset of 100 V, and maintained at 20 °C. The scan time was set to 1 scan per second. Calibration is performed using a standard cesium iodide (CsI) solution. Sample illumination was carried out by irradiation of the nano electrospray emitters with a 455 nm LED (M455L4, *Thorlabs*) at 30-50 µmol m^−2^ s^−1^ (Extended Data Fig. 6a), using an illumination setup similar to that described by Camacho *et al*. (2019).^47^ Acquisition and data processing are achieved using MassLynx™ V4.1 SCN901. Prior to measurements, CRY1 was buffer-exchanged in 200 mM ammonium acetate, 10 mM DTT and pH 7.4, using Zeba Micro Spinbuffer exchangers (7 kDa MWCO, *Thermo Fisher Scientific*).

### nESI-MS based rate constant determination

The measurements of the rate constants were performed following the same procedure as for the nESI-MS measurements except for the scan time, which was set to 10 scans per second. The intensity (counts) of each species (monomer, dimer, tetramer) during the measurement period was estimated by generating an extracted ion chromatogram using MassLynx™ V4.1 SCN901. Initial concentrations of each species (monomer, dimer, tetramer) were determined from the measured signal intensities, assuming a total sample concentration of 15 µM relative to the molecular mass of the monomer. Kinetic data were analyzed by using DynaFit^48^ software and a reversible two-step kinetic model (Fig. 1l) based on the differential equations 1-3.

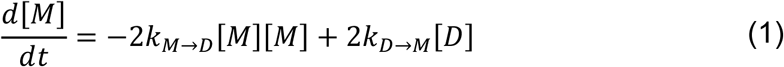

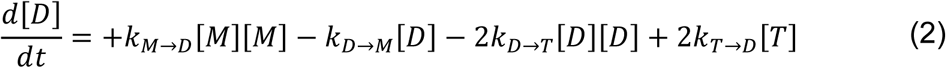

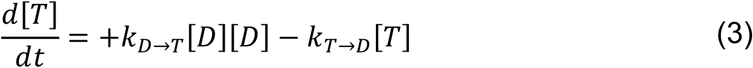

### LILBID-MS

Prior to analysis by LILBID-MS^38^, CRY1 and BIC1^3CA^ were buffer-exchanged as described for nESI-MS. All measurements were performed at 20 °C, using 4 μL of buffer-exchanged and degassed sample for each analysis. Microdroplets with a diameter of 50 μm are generated using a piezo-driven droplet generator (MD-K-130, *Microdrop Technologies GmbH,* Norderstedt, Germany) operating at 10 Hz. The droplets are transferred into a vacuum of about 10^-5^ mbar and irradiated with a pulsed infrared laser (*Innolas Spitlight 400, Continuum PowerLite 8000*) tuned to the vibrational absorption maximum of water (2.8 μm). Absorption of laser energy (10-23 mJ per pulse) by water molecules causes an explosive expansion of the droplets, resulting in the release of analyte ions into the gas phase. The ions are accelerated using a pulsed electric field and analysed with a home-built time-of-flight mass spectrometer in negative ion mode. Each mass spectrum represents an average signal from 1,000 individual microdroplets. Mass spectra are calibrated against a 10 μM aqueous bovine serum albumin standard and subsequently processed by smoothing and background subtraction. The sample was exposed to blue light (455 nm LED, M455L4, *Thorlabs*) at 100 µmol m^−2^ s^−1^ inside the droplet generator via a coupled optical fibre that was installed on the generator (Extended Data Fig. 6b).

For qLILBID-MS^39,40^ experiments the laser energy transferred into the droplets is correlated to the degree of complex dissociation. Therefore, the explosion of the irradiated droplets is visualized by illuminating the explosion with a green laser (Minilite I, *Continuum*, San Jose, USA) and the resulting images are captured using a high-speed camera (DFK 23UP031, *Imaging Source*, Bremen, Germany). Each individual droplet yields both a mass spectrum and a corresponding image of the explosion. The explosion width for each droplet is determined by analysing the recorded images with OpenCV, using a custom written Python script. The resulting dissociation plots allow K_D_ determination.

### TEM negative staining

Carbon coated copper grids (400 mesh) were hydrophilized by glow discharging (*PELCO easiGlow*, Ted Pella, USA). 5 µl of a protein suspension with a concentration of 30 µg/ml was applied onto the hydrophilized grids and stained with 2% uranyl acetate after a short washing step with filtered H_2_O_bidest_. If applicable, samples were prepared under blue (465 ± 5 nm, 115 µmol m^−2^s^−1^) or safe red light, respectively. Samples were analysed with a JEOL JEM-2100 transmission electron microscope using an acceleration voltage of 120 kV. For image acquisition a F214 FastScan CCD camera (*TVIPS, Gauting*) was used.

## Data Availability

All data generated in this study are provided as Source Data with this paper. Gene accessions codes are At3G52740 (*BIC1*) and At4G08920 (*CRY1*).

## Acknowledgements

We thank Konstantinos Stamatakis for his support and fruitful discussions during the kinetic modelling process, and Josef Wachtveitl for providing access to the absorption spectrometer. We also acknowledge Jeanette Schermuly for technical assistance and Stephan Kiontke for his contributions to the recombinant expression of BIC1 and for helpful discussions. Funding was provided by the German Research Foundation (DFG): A.J. was supported through the Collaborative Research Center CRC 1507 “Membrane-Associated Protein Assemblies, Machineries and Supercomplexes”, N.M. through DFG Grant 557111829, L.-O.E. through DFG grant ES152/18 and A.B. through DFG grant BA985/15-1.

## Author contributions

A.J., N.M., L.-O.E., A.B. designed and coordinated the experiments. P.G., A.J., N.N. and D.K. expressed and purified the proteins. A.J. performed mass spectrometry, UV-Vis spectroscopy and kinetic modelling. N.N. and T.H. acquired and interpreted the TEM data. N.N. acquired and interpreted the SEC data. A.J. wrote the original draft. A.J., N.M., A.B., P.G., L.-O.E. edited the manuscript.

