## Extended Data for "Mechanistic insight into the oligomerisation of *Arabidopsis* CRY1 and its inhibition by BIC1"

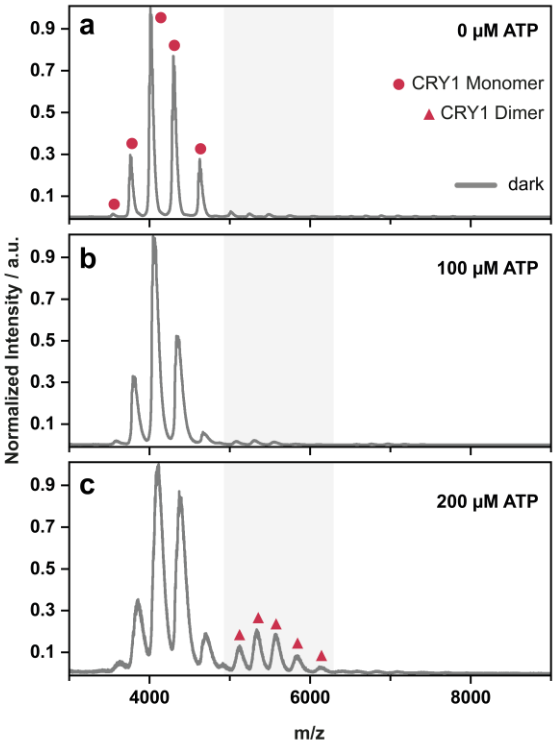


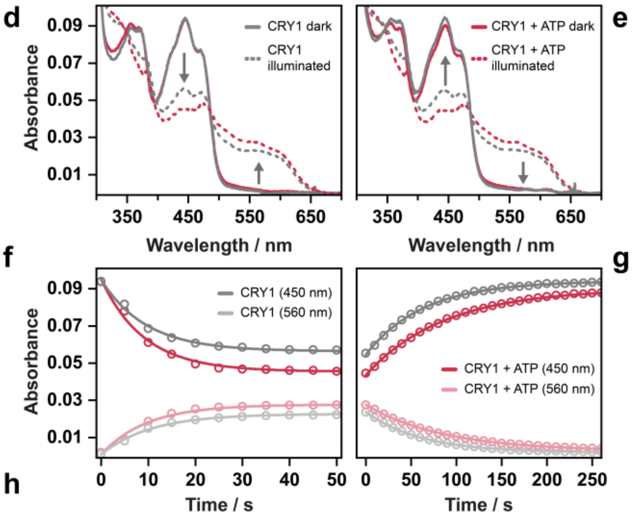


h

| | Photoreduction | | Thermal decay | |
|---|---|---|---|---|
| | $t_{1/2\ (450\ nm)}$ / s | $t_{1/2\ (560\ nm)}$ / s | $t_{1/2\ (450\ nm)}$ / s | $t_{1/2\ (560\ nm)}$ / s |
| **- ATP** | 7.3 ± 0.9 | 7.275 ± 0.005 | 42 ± 4 | 42.5 ± 0.5 |
| **+ ATP** | 6.44 ± 0.15 | 6.78 ± 0.03 | 67 ± 9 | 65 ± 8 |

**Extended Data Fig. 1: ATP modulates CRY1 dimerization in darkness and enhances FAD photoreduction kinetics. a-c**, nESI-MS spectra of CRY1 in the absence of blue light at different ATP concentrations. Incubation with 200 µM ATP resulted in an increase of dimeric CRY1 species (**c**) compared to samples with 0 (**a**) or 100 µM ATP (**b**). **d-h**, ATP accelerates photoreduction and stabilizes the reduced state of the FAD cofactor. Addition of 100 µM ATP enhanced the rate of $FAD_{ox}$ to FADH° photoreduction by ~12% (**f, h**) and slowed the thermal reoxidation of FADH° to $FAD_{ox}$ by ~60% (**g, h**).

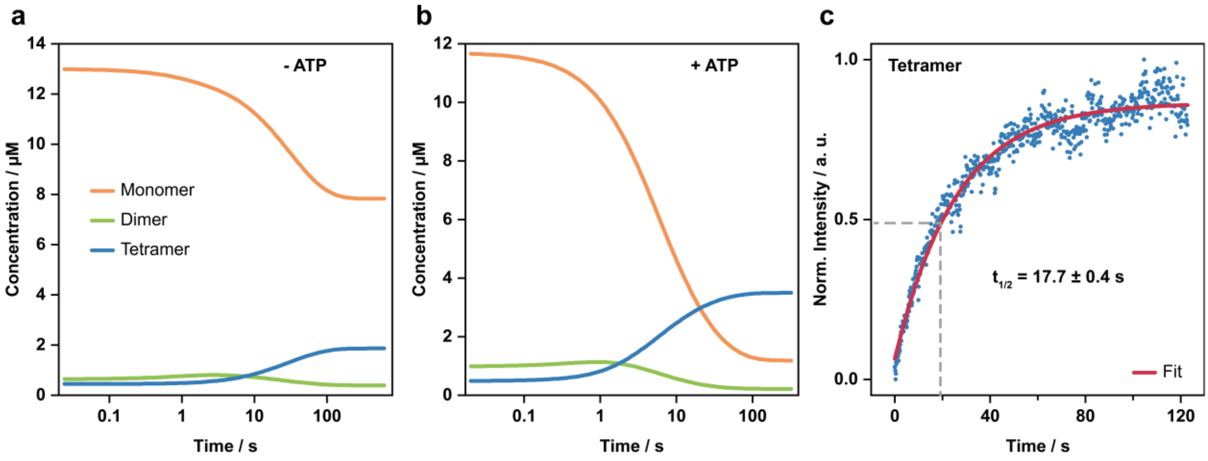


**Extended Data Fig. 2: Light-dependent oligomerisation dynamics of CRY1 analysed through kinetic modelling and time-resolved mass spectrometry.** Concentration profile of CRY1 monomer, dimer, and tetramer in absence (**a**) and presence of 100 µM ATP (**b**) during exposure to blue light obtained from the fit of the kinetic model. **c**, Fitting the tetramer formation data obtained from time-resolved nESI-MS measurements using a single-exponential model, yielding a half-life of $t_{1/2}$ = 17.7 ± 0.4 seconds.

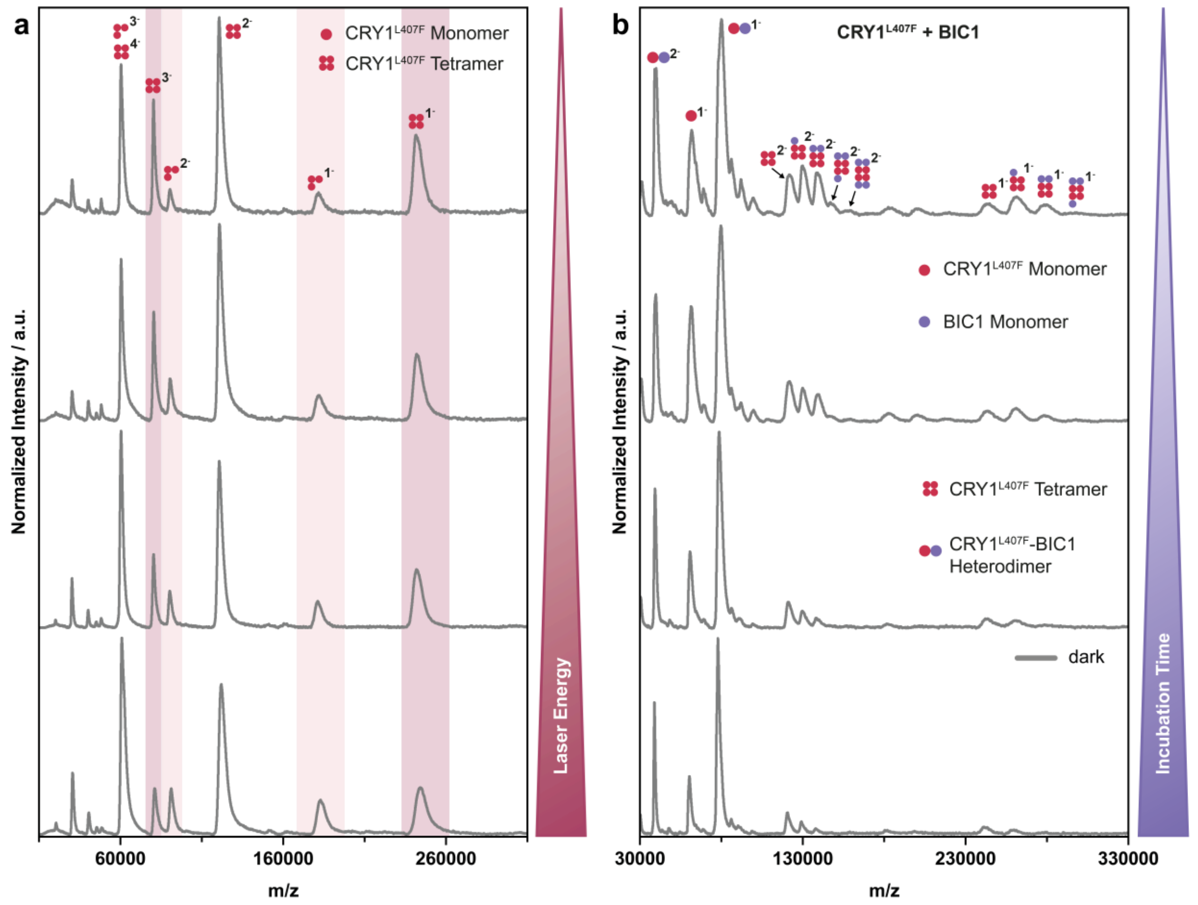


**Extended Data Fig. 3: LILBID-MS spectra showing dissociation of the CRY1 tetramer into a trimer upon increasing laser energy and time-dependent disassembly of CRY1$^{L407F}$ tetramers by BIC1**. **a**, Increasing laser energy results in a decrease of the CRY1 tetramer signal (highlighted in dark red) and the appearance of a trimeric species (highlighted in light red), indicating partial dissociation of the complex. **b**, BIC1 induces a progressive disassembly of CRY1$^{L407F}$ tetramers in darkness with increasing incubation time by binding to individual CRY1$^{L407F}$ protomers.

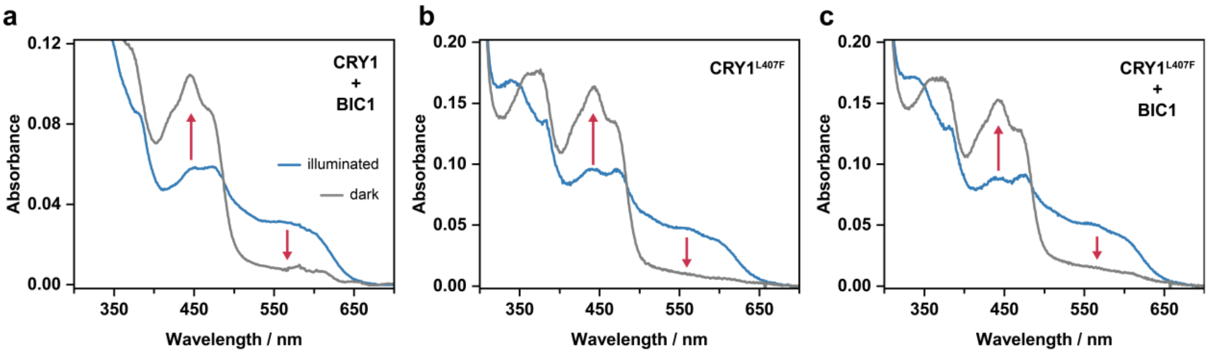


**Extended Data Fig. 4: Absorption spectra slowing the thermal reoxidation of the FADH° semiquinone to $FAD_{ox}$ in wild-type CRY1 and CRY1$^{L407F}$ mutant incubated with BIC1. a**, Thermal reoxidation of the reduced FAD chromophore of CRY1 incubated with BIC1. **b, c**, Thermal reoxidation of the reduced FAD chromophore of CRY1$^{L407F}$ showing that the L to F replacement at position 407 incubation with BIC1 in an equimolar ratio does not impair the chromophore's photoreduction.

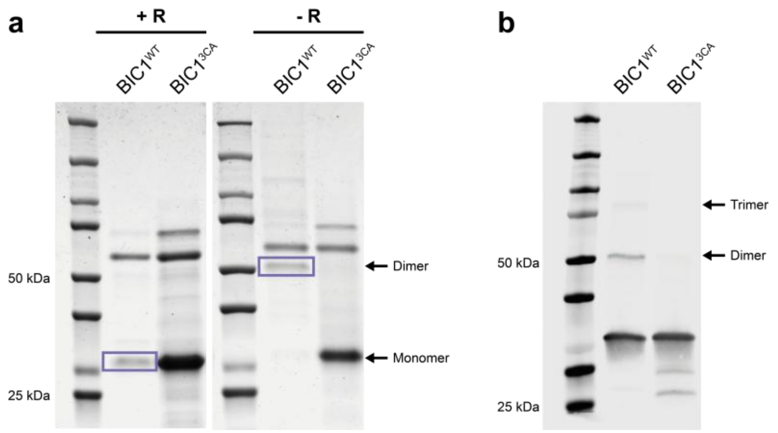


**Extended Data Fig. 5 Wild-type BIC1 but not the BIC1$^{3CA}$ triple mutant forms intermolecular disulfide bridges. a**, Coomassie-stained SDS-PAGE gel with 3 µg of protein loaded in each lane of wild-type (BIC1$^{WT}$) or the C to A triple mutant (BIC1$^{3CA}$) in presence (+R) or absence (-R) of 10 mM reductant Tris(2-carboxyethyl)phosphine-hydrochloride – TCEP in SDS loading buffer, boiled for 10 min. The BIC1 monomers run at about 25 kDa. Black arrows indicate monomers, dimers or trimers of BIC1. **b**, Western blot of SDS-PAGE gel with 1.5 µg of protein loaded per lane and probed with αHis antibody showing that, in contrast to BIC$^{WT}$, the BIC$^{3CA}$ triple mutant does not form oligomers.

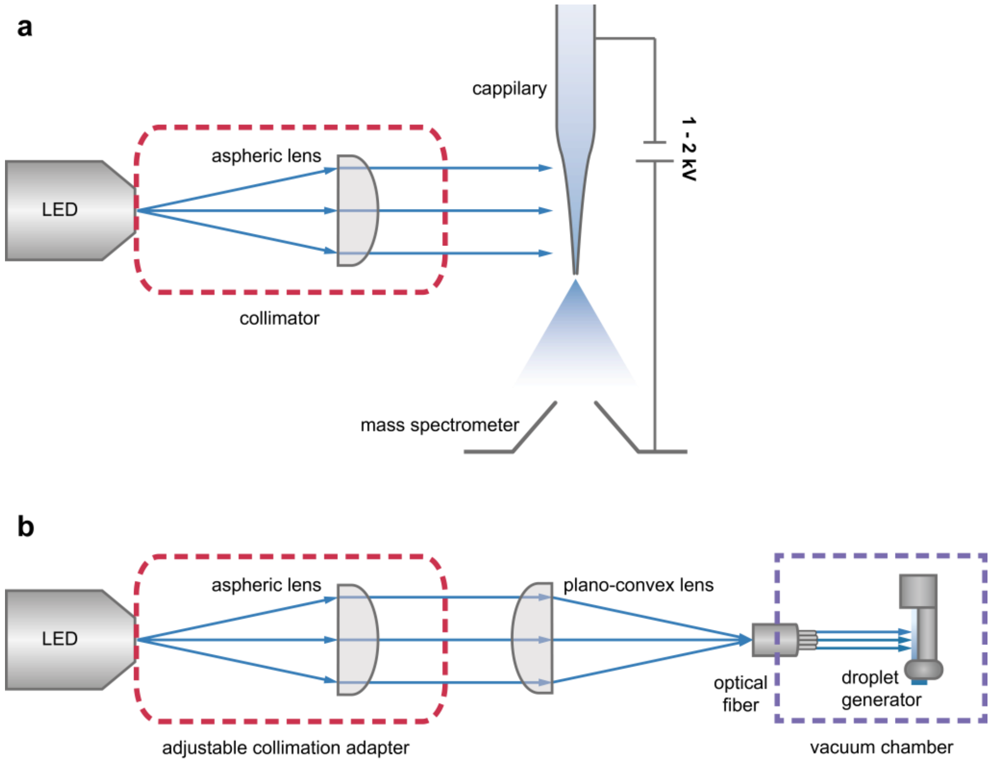


**Extended Data Fig. 6: LED-based illumination setups for LILBID-MS and nESI-MS. a**, Schematic representation of the LED-based illumination setup for nESI-MS. A 445 nm LED is directed onto the sample-filled capillary via a collimator, allowing illumination of the electrospray emitter at the front end of the mass spectrometer. **b**, Schematic representation of the integrated light source used in the LILBID mass spectrometer. Light from a

455 nm LED is coupled into an optical fibre via an adjustable collimator and lens system and directed onto the droplet generator within the vacuum chamber.
